# Single-cell study of the FCG rat model and cross-species comparison reveal stronger gene regulation by sex chromosome complement in the medial septum

**DOI:** 10.64898/2026.11.19.767942

**Authors:** Victoria Turnbill, Guanglin Zhang, Erin Harris, Juliana Shin, Xuqi Chen, In Sook Ahn, Megan Rivera, Ethan Tran, Dhruvikaa Reddy, Rehan Wani, Melinda Dwinell, Vincent Harley, Aron M. Geurts, Arthur P. Arnold, Debra A. Bangasser, Xia Yang

**Affiliations:** Department of Integrative Biology and Physiology, University of California, Los Angeles, CA, USA; Neuroscience Interdepartmental Program (NSIDP), University of California, Los Angeles, CA, USA; Neuroscience Institute, Georgia State Univ., Atlanta, GA, USA; Center for Behavioral Neuroscience, Atlanta, GA, USA; Molecular and Medical Pharmacology, University of California, Los Angeles, CA, USA; Hudson Institute of Medial Research, Melbourne, VIC, Australia; Department of Physiology, Medical College of Wisconsin, USA

## Abstract

Sex differences are prominent in neuropsychiatric, neurological, and neurodegenerative conditions. While both gonads (ovaries vs. testes) and sex chromosome complement (SCC; XX vs. XY) contribute to sex differences, these effects are hard to differentiate in humans and gonadal contributions have been considered dominant in the brain. The four core genotypes (FCG) mouse model distinguishes between SCC and gonadal effects but whether findings are conserved across species remains unclear. Here, we present the first application of the novel FCG-derived rat model to explore cell-type-specific sex differences in the medial septum (MS), which shows robust structural sex differences, and convergence/divergence between species. Differential gene expression, pathway, transcription factor, and gene regulatory network analyses show dominant SCC effects and concordant gonadal effects on gene and network regulation in rat MS cell types. Comparison between the FCG rat and mouse models find convergence across species on the dominance of the SCC in regulating gene expression in concert with gonadal effects in the MS, despite divergence in individual sex-biased genes. In both species, the sex-biased, cell-type-specific MS genes and networks are associated with diverse diseases relevant to MS functions, such as anxiety, Alzheimer’s Disease, epilepsy, and schizophrenia, that can be targeted by proposed therapeutic candidates. Our cross-species investigation identifies a conserved dominance of SCC effects in sex-biased gene expression in the MS, thereby challenging the classical gonad-centric view of sexual differentiation in the brain and highlighting the MS as an important region for the study of sex differences driven by the sex chromosome complement.

## Introduction

Sex differences are found at all levels of human biology with notable differences in the risk, prevalence, symptomology, and progression of various neuropsychiatric, neurological, and neurodegenerative conditions. For example, some neuropsychiatric conditions with origins in development – such as Autism Spectrum Disorder (ASD), Attention Deficit Hyperactivity Disorder (ADHD), and schizophrenia (Scz) – occur several times more often in males than females (McCarthy et al. 2017). The opposite is true in some adult-onset conditions, including Major Depressive Disorder (MDD), bipolar disorder, generalized anxiety, panic disorder, and anorexia nervosa (McCarthy et al. 2017). Neurodegenerative diseases are also sex-biased: Multiple Sclerosis and Alzheimer’s Disease (AD) affect roughly two to three times as many women as men while Parkinson’s Disease (PD) and Amyotrophic Lateral Sclerosis (ALS) affect up to 3 times as many men as women (McCarthy et al. 2017). Understanding the underlying sexually biased mechanisms of development and disease is key for the development of effective treatments and preventions (Ngun et al. 2011; McCarthy et al. 2017); however, this remains under-studied (Arnold et al. 2023).

Sex differences in brain region- and cell-type specific gene networks arise from a combination of sex chromosomal and gonadal factors, which are difficult to dissect in typically developing males and females due to their overlapping effects, so special models are required to pinpoint specific regulators (Arnold 2017; Arnold 2020; Arnold et al. 2026). Previous work, including ours, has begun to address this gap by using the Four Core Genotypes (FCG) mouse models that are genetically modified to uncouple gonadal (ovaries vs. testes) from sex chromosome complement (SCC; XX vs. XY) effects. Applying high throughput molecular profiling to these mouse models helps partition sex-biasing factors in the brain and facilitates the understanding of sex-biased human brain disorders (Blencowe et al. 2022; Shin et al. 2025). As of 2021, over 80 publications had used the FCG mouse model of sex differences to study everything from metabolism and hypertension to behavior and AD (Arnold 2020). However, mouse models do not capture many aspects of human pathophysiology, and additional models are needed to facilitate translational and generalizable discoveries (Ellenbroek and Youn 2016; Hickman et al. 2017; Guma et al. 2024). For these reasons, Arnold et al. recently developed an FCG-derived rat model of sex differences to complement the existing FCG mouse model and enable cross-species comparisons (Arnold et al. 2026). In this new model, the *Sry* genes responsible for testis development are either removed from the genome entirely (resulting in ovary development) or translocated from the Y chromosome into an autosomal location (Arnold 2020; Arnold et al. 2026). Gonad development and the SCC are functionally separated: XY rats can be produced with testes (XYT), and XX rats can have either testes (XXT) or ovaries (XXO). Thus, use of the FCG-derived rat model can determine whether typical sex differences (XYT vs. XXO) are the result of gonadal (XXT vs. XXO) and/or SCC (XYT vs. XXT) effects and/or their interactions – which together drive sex differences (Arnold 2009; Arnold 2020; Arnold et al. 2023). This study is the first biological application of the FCG-derived rat model to characterize sex-biasing factors and regulators of sex differences in the rat brain.

Among brain regions, the medial septum (MS) is an important yet understudied locus for examining sex differences. The MS is a key regulatory hub in spatial navigation, learning, memory consolidation and integration, hippocampal theta rhythm generation, wakefulness, and social memory (Muller and Remy 2018; Griguoli and Pimpinella 2022; Leung and Ma 2022; Song et al. 2025). It is implicated in sexually biased human conditions including epilepsy, Dravet and Rett syndromes, Scz, post-traumatic stress disorder (PTSD), AD, ASD, stress, anxiety, and pain perception (Ma and Leung 2014; McCarthy et al. 2017; Leung and Ma 2018; Muller and Remy 2018; Griguoli and Pimpinella 2022; Leung and Ma 2022; Fan et al. 2023; Wu et al. 2024; Song et al. 2025; Wu et al. 2025; Zhu et al. 2025). Limited existing data on the MS show interesting sex differences in this brain region. Mouse studies have shown that it is larger in females than males – a phenotype that is likely caused by SCC effects since removal of gonadal hormones before puberty did not alter the female > male relationship (Corre et al. 2016; Vousden et al. 2018; Raznahan and Disteche 2021). Additionally, peak neurogenesis of MS cholinergic neurons during development occurred earlier in male mice than in females (Schaevitz and Berger-Sweeney 2005). Greater vulnerability to impaired, MS-mediated memory formation has also been shown in males compared to females as a result of corticotropin releasing factor (CRF), and memory formation was independent of circulating ovarian hormones (Wiersielis et al. 2019). Finally, our recent work in FCG mice has uncovered widespread sex-biased differences in gene expression across cell types in the MS which are explained by substantial SCC effects as compared to gonadal hormone effects (Shin et al. 2025). Notably, sex chromosome-associated differentially expressed genes (DEGs) in the FCG mouse model were enriched for various human disorders ranging from ADHD and Scz to insomnia and metabolic disorders (Shin et al. 2025). In sum, these findings highlight the MS as an important region for the study of sex differences related to the SCC and numerous human pathologies. However, whether these findings hold true in other species is unknown.

Here, using the new FCG-derived rat model and single nuclei RNA sequencing (snRNAseq), we identify sexually dimorphic DEGs, pathways, and functional networks in the rat MS and further dissect these differences into their putative underlying sex-biasing mechanisms (i.e., gonads vs SCC). We then integrate our results from the FCG-derived rats with a broad screening of ∼200 human GWAS datasets relevant to brain disease and available at GWAS catalog to reveal associations of the sex-biased cellular and molecular networks in the MS with specific human brain disorders (Cerezo et al. 2025). Then, we compare our results with those from the FCG mouse to make cross-species comparisons and determine which human pathologies may be preferentially studied in either the FCG-derived rat or FCG mouse models to facilitate future translational research. Finally, we propose drug repositioning candidates as potential therapeutics for sex-biased, MS-linked human conditions. and sex factor. Significance was set to FDR < 5%.

## Results

### snRNA-seq reveals cellular landscape of the MS in FCG rats

Cells across all 3 experimental groups (n = 9 pools, n=3 pools/group, and 2 rats/pool for a total of 18 rats) were clustered based on transcriptomic similarity and projected to 2D space using UMAP. After quality control and filtering, 70,512 nuclei were included in our analysis: 25,803 XYT nuclei, 25,379 XXT, and 19,339 XXO (**Sup. Fig. 2A**). Using the Allen Institute’s online tool MapMyCells, we identified 10 cell types: mature and immature oligodendrocytes, oligodendrocyte precursor cells (OPCs), endothelial cells, vascular cells, microglia, two subtypes of GABAergic neurons (1 and 2), glutamatergic neurons, and cholinergic neurons (**Sup. Fig. 2B**). We verified cell type annotations by cell-type-specific marker expression (**Sup. Fig. 2C**) and used R package MetaCell (Baran et al. 2019; Bilous et al. 2022) to construct 5,795 metacells (to reduce data sparsity and mitigate inflation of statistical power) parsed by sample and cell type (**Sup. Fig. 2D-F**).

### Sex-biasing factors do not significantly affect MS cell type proportions

We used R package speckle (Phipson et al. 2022) to perform cell type proportion analysis on raw nuclei counts and determine whether the proportion of each cell type differs across experimental groups (**Sup. Fig. 2G-H**). No significant group differences were detected – suggesting that sex-biasing factors do not directly modify relative composition of individual cell types in the rat MS.

### DEGs Reveal Stronger SCC than Gonadal Effects with Concordance between the Two

We performed DEG analysis using Limma(Ritchie et al. 2015) at FDR <5% across three pairwise comparisons (Typical: XYT vs. XXO; Gonadal: XXT vs. XXO; SCC: XYT vs. XXT). The full DEG list is available in **Sup. Tab. 3.** As a positive control, we first examined the X- and Y-linked genes expected to be sex-biased. In Typical and SCC comparisons, we observed the expected higher expression in XX than XY of certain X-linked genes (e.g., *Xist*) and the expected higher expression in XY than XX of certain Y-linked genes (e.g., *Uty*; **Fig. 1A**). For each sex-biasing effect, astrocytes had more DEGs than the other cell types (**Fig. 1B**) and we also identified roughly equal numbers of up-vs. down-regulated DEGs (**Fig. 1C**) and the expected proportions of sex-linked vs. autosomal DEGs (**Fig. 1D**). We next parsed the total DEGs into those found across 2 or more sex factors (Shared) and those identified only within one sex factor: Typical, Gonadal, or SCC (**Fig. 1E**). Agreeing with the dominant SCC effects on MS volume and gene expression in mouse MRI and transcriptomic studies (Corre et al. 2016; Raznahan and Disteche 2021; Shin et al. 2025), we identified a larger contribution of SCC-unique DEGs (3,777) compared to Gonadal-unique DEGs (2,254). There were 5,169 Typical-unique DEGs and 7,614 Shared DEGs between sex factors.

**Figure 1.**
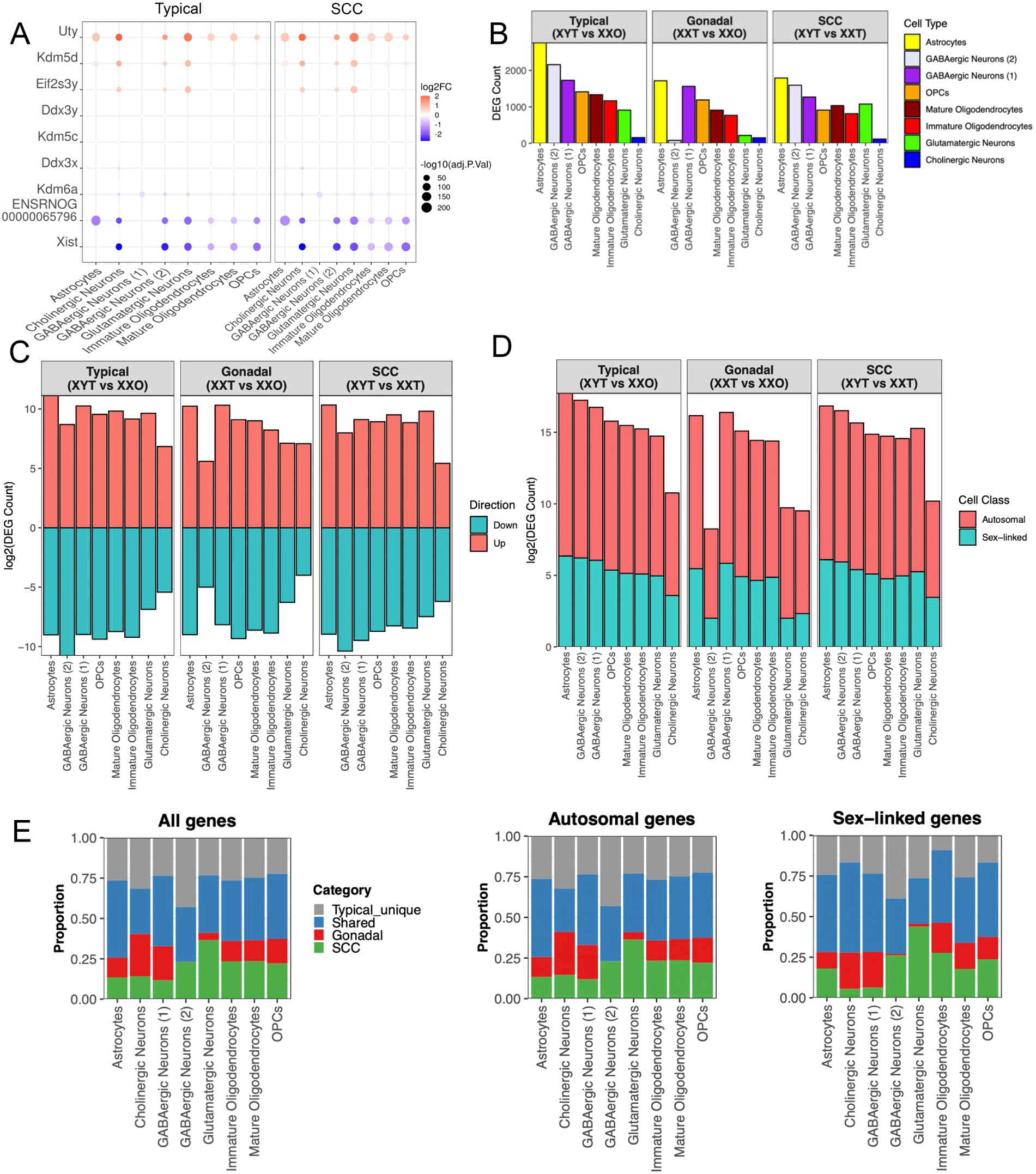
Cell-type-specific DEGs across experimental groups for the various sex factors in the FCG-derived rat MS. (A) Differential expression of sex-linked genes by cell type in the Typical comparison (XYT vs XXO) and the SCC comparison (XYT vs. XXT) reveal expected higher expression of certain Y-linked genes in males (red) and higher expression of certain X-linked genes in females (blue). (B) Numbers of DEGs per cell type for typical, gonadal, and SCC effects. (C) Numbers of “Up” (XYT > XXO, XXT > XXO) vs. “Down” (XYT < XXO, XXT < XXO, and XYT < XXT) regulated DEGs for Typical, Gonadal, and SCC sex differences per cell type. (D) The proportions of sex-linked vs autosomal DEGs from the Typical, Gonadal, and SCC comparisons per cell type. (E) Bar plots indicating the Typical, Gonadal, and SCC DEG set overlap, where DEGs are classified as being Typical-unique, Gonadal-unique, SCC-unique, or Shared (being present in at least two DEG sets), show greater numbers of SCC-unique DEGs than Gonadal-unique in the majority of cell types. Results are separated based on genomic location of the genes (i.e. All Genes, Autosomal, or Sex-Linked). Together, these analyses indicate that SCC contributes more than gonadal sex to transcriptional differences in the rat MS.

Our prior study found the expression of corticotropin releasing factor (CRF)-binding protein (encoded by *Crhbp*), which sequesters CRF to reduce its bioavailability, was higher in bulk MS punches in female than male rats(Wiersielis et al. 2019). This effect was associated with a CRF-biased memory impairment in male but not female rats that was independent of ovarian hormones – suggesting an SCC effect. The current snRNAseq results reveal that XXT and XXO groups both express more *Crhbp* than XYT in GABAergic neurons in the MS (**Sup. Fig. 3**). These data indicate that the sex difference observed in bulk MS tissue reflects cell type-specific expression within GABAergic populations. Moreover, elevated *Crhbp* expression in both XX genotypes with different gonads implicates SCC, rather than gonadal hormones, as the primary driver of this effect, consistent with our initial hypothesis(Wiersielis et al. 2019).

We then performed a Pearson correlation analysis of DEGs across cell types and sex-biasing factors to determine global patterns of similarity in gene expression changes. This resulted in DEG clusters representing similar gene expression changes (**Fig. 2A**). Except for the Gonadal Neurons cluster that contains gonadal effect DEGs across multiple neuronal subtypes, all clusters were homogeneous for cell type – indicating that patterns in gene expression changes across each sex factor were cell-type specific. Also, in every cell type but OPCs, Typical and SCC DEGs clustered together while Gonadal DEGs clustered independently – indicating a stronger similarity between Typical and SCC effects on gene expression patterns than between Typical and Gonadal effects. This suggests that SCC effects, rather than gonadal effects, likely drive the majority of gene expression changes in the Typical sex differences.

Finally, most correlations are positive in **Fig. 2**, suggesting that gonadal and SCC effects induced changes in the DEGs in the same direction to contribute synergistically to gene expression changes observed in the Typical sex differences.

### Pathway and Transcription Factor Enrichment Further Shows Gonadal and SCC Concordance

Next, we performed pathway enrichment analysis for each of the DEG clusters (**Fig. 2B, Sup. Tab. 4**). Top enriched pathways for both SCC and gonadal DEGs relate to RNA splicing, protein transport, or synapse structure and function. These pathways show higher expression in male gonad or XY genotype (XYT>XXO, XYT>XXT, XXT>XXO) in most cell types except the Typical and SCC GABAergic Neurons (2) cluster. These results indicate pathway level concordance between gonadal and SCC effects which together contribute to sex difference.

**Figure 2.**
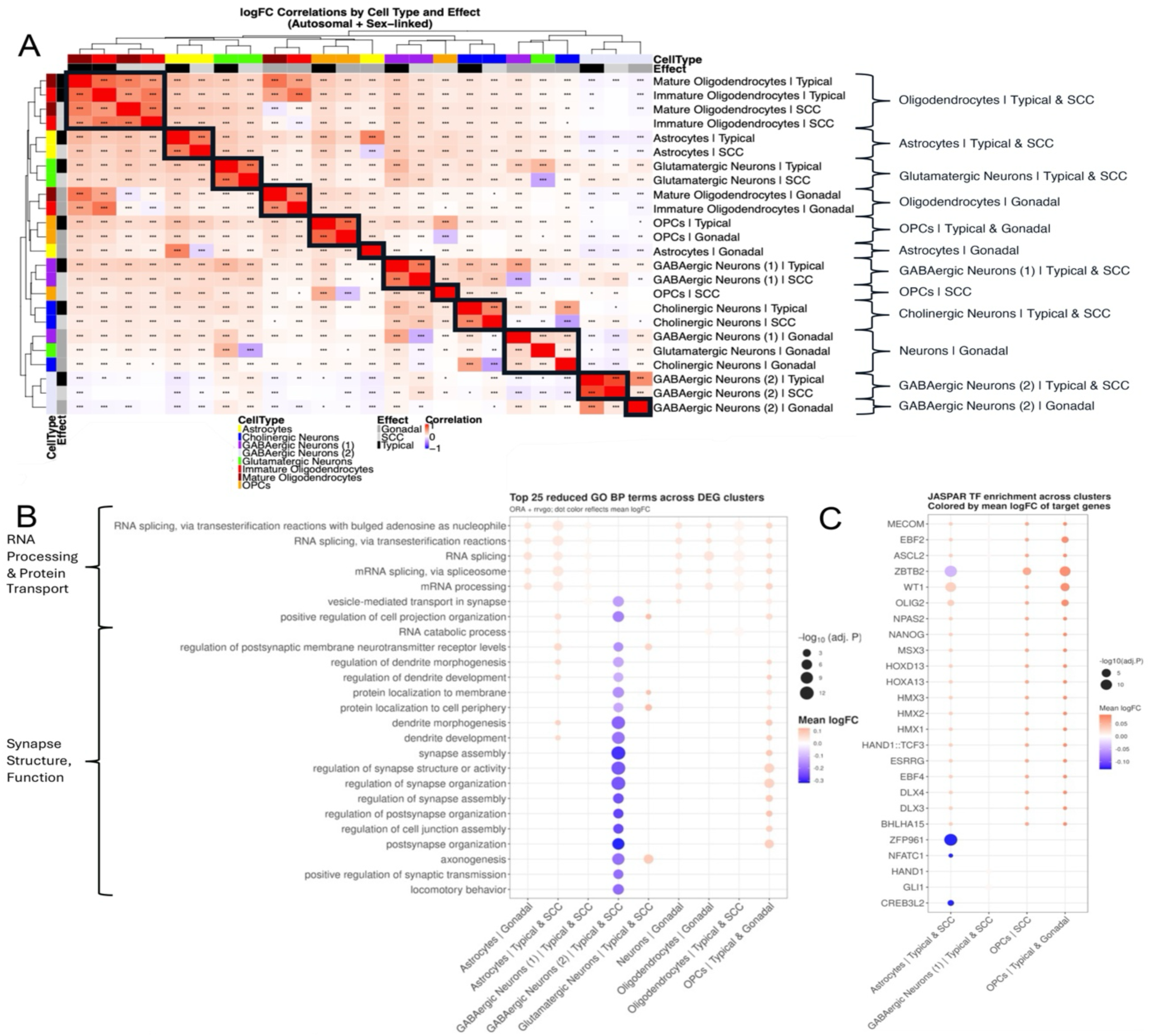
DEG clustering and pathway and transcription factor enrichment show stronger similarity of the SCC effects to Typical sex differences and concordance between SCC and Gonadal effects. (A) Heatmap shows pairwise correlations of normative log2FC values across cell types and sex effects, and dendrograms reveal hierarchical clustering of cell type and sex effect DEG sets where SCC and Typical DEG sets cluster more closely together and Gonadal DEG sets cluster more independently and nearly all correlations are positive. (B) rrvgo reduction of overrepresentation analysis (ORA) of the identified DEG clusters against GO:BP gene sets shows mostly higher expression in males of pathways based on DEG expression related to RNA processing and protein transport and synapse functioning as well as concordant SCC and gonadal effects in most cell types. (C) rrvgo reduction of ORA against target genes of transcription factors from the JASPAR PWM Mouse 2025 database shows that more SCC DEG clusters are enriched for transcription factor targets and are mostly upregulated in males, with concordant SCC and gonadal effects for OPCs. These results demonstrate that the SCC DEG clusters contribute more to the Typical sex differences than gonadal clusters in terms of pathway and transcription factor regulation and that SCC and gonadal effects are concordant.

To identify potential regulators of the DEGs, we performed transcription factor (TF) enrichment analysis on the clusters (**Fig. 2C, Sup. Tab. 5**). With few exceptions, we found more SCC DEG clusters showing TF enrichment, higher expression of the target genes of the enriched TFs in male genotypes relative to females, and concordance between gonadal and SCC effects. Most TFs (e.g., MECOM, ASCL2, ZBTB2, NANOG, and OLIG2) are implicated in various functions related to development, cell differentiation, and cell proliferation(Nagao et al. 2016; Liu et al. 2019; Jung et al. 2022; Fang and Bai 2023; Gulla et al. 2026). However, DEG targets of both ESRRG and NPAS2 are expressed higher in males than females in SCC and Gonadal effects in astrocytes and OPCs. These two TFs are related to hormone signaling pathways and molecular clock mechanisms respectively (Giguère 2002; Peng et al. 2021; Fox et al. 2025). The importance of *NPAS2* in time-keeping functions supports the known function of the MS as a center of rhythmogenesis throughout the basal forebrain and in the septo-hippocampal circuit (Muller and Remy 2018; Song et al. 2025) and suggests that rhythmogenesis is subject to sex-biased regulation by both SCC and gonadal effects.

### Gene Networks Are Regulated by Both SCC and Gonadal Effects

To further identify potential non-TF regulators of the DEGs, we used SCING to perform global network inference within cell types and Mergeomics wKDA to identify key drivers whose neighborhood subnetworks were significantly enriched for DEGs for each sex factor (Typical, Gonadal, and SCC) (Shu et al. 2016; Ding et al. 2021; Littman et al. 2023). While some cell types displayed gene regulatory networks (GRNs) that were largely enriched for DEGs specific to SCC effects, as expected, other cell types showed strong enrichment for DEGs specific to gonadal effects as well. For instance, Cholinergic Neurons and GABAergic Neurons (2) mainly show SCC effect GRN regulation (**Fig. 3A-B**); the rest show both SCC and Gonadal regulation (**Fig. 3C-H**). Some GRNs are regulated almost solely by gonadal effects (e.g. one main subnetwork in Astrocytes, **Fig. 3C**). Thus, while earlier results showed strong SCC influences on gene, pathway, and TF expression, these GRN results suggest that both SCC and gonadal effects play important roles in regulating expression of functional subnetworks in a cell-type specific way.

**Figure 3.**
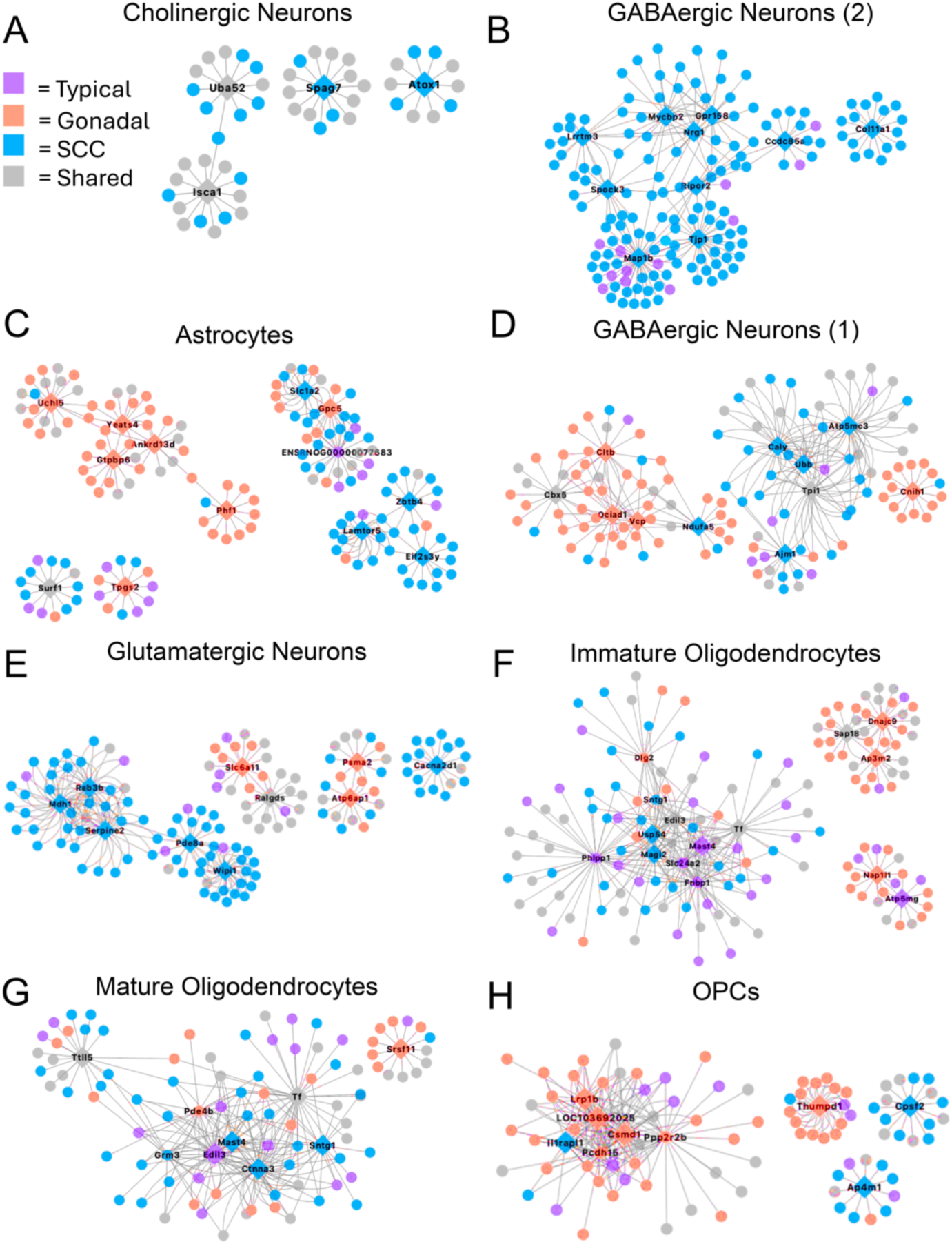
Gene regulatory network (GRN) modeling reveals functional networks and key drivers of sex differences within cell types in the MS regulated by both SCC and gonadal effects. Visualization of GRNs by cell type around key drivers identified for sex-factor-associated DEGs affected by both SCC and gonadal effects. Key drivers are shown as nodes in diamond shapes, subnetwork member genes are denoted by smaller circular nodes, and all shapes are colored by DEG representation within Typical (purple), SCC (blue), Gonadal (red), or Shared (gray) sex factors. GRNs and key drivers within Cholinergic Neurons (A) and GABAergic Neurons (2) (B) are regulated by SCC effects while Astrocytes (C), GABAergic Neurons (1) (D), Glutamatergic Neurons (E), Immature Oligodendrocytes (F), Mature Oligodendrocytes (G), and OPCs (H) are all regulated by both SCC and gonadal effects. Gene regulatory network analysis reveals that cell-type-specific regulatory networks are jointly shaped by SCC and gonadal effects, with SCC dominant in select cell types.

Network key drivers are listed by cell type in **Sup. Tab. 6** with their known functions. Key drivers are not shared across neuronal cell types, but there is some overlap in the potential functions of their GRNs. OPCs do not share key drivers with either mature or immature oligodendrocytes, however several are shared between immature and mature oligodendrocytes – indicating that there is more functional similarity between oligodendrocyte subtypes. One astrocytic key driver, *Rora*, has been shown previously to regulate various genes involved in circadian rhythm (Sato et al. 2004). This is especially relevant to the previously known function of the MS in wakefulness, anesthesia response, and maintenance of the sleep-wake cycle and potentially implicates astrocytes as timekeeping cells within the MS (Leung and Ma 2022; Song et al. 2025). **Sup. Tab. 6** also includes a list of diseases known to be associated with key drivers which provides further evidence that the MS may be important for the pathogenesis of certain conditions with sex biases linked to this brain region previously, including epilepsy, psychosis, Scz, and AD.

### PharmOmics analysis proposes sex-specific therapeutics acting on GRNs across cell types

We next used PharmOmics for drug repositioning analysis to propose candidate drugs predicted to act on sex-biased GRNs across multiple cell types in the MS (**Sup. Tab. 7**). The 25 drugs with the highest overlap with network key driver genes across cell types are listed in **Sup. Tab. 8**. We hypothesize that, by acting on many key drivers for sex-biased GRNs across multiple cell types in the MS, these drugs will have greater efficacy than direct or indirect manipulations of one or a few key genes in one sex versus the other.

Among the top candidates listed in **Sup. Tab. 8**, there are four drugs that interact with gonadal steroid hormone receptors: β-estradiol, Ethinylestradiol, Flutamide, and Diethylstilbestrol. Methyltestosterone is also among the significant drugs predicted, however it was not among the top 25 ranked candidates. This provides additional support for the important role of gonadal hormones in regulating these GRNs across cell types in the MS. Other drug classes may act on either gonadal hormones or the SCC since both gonadal and SCC effects were shown to regulate these key GRNs. Additionally, of the 25 top drug candidates, at least 16 have been shown to have sex specific effects, side effects, pharmacokinetics, or outcomes (**Sup. Tab. 8**). This further supports the idea that the GRNs constructed in **section 2.5** are regulated in sex-biased ways and signifies the importance of patient sex for designing treatment protocols. Finally, most drugs pass the blood brain barrier and thus are intriguing future therapeutic candidates for treatment of MS-linked conditions (**Sup. Tab. 8**).

### Most differentially expressed macrogenes (DEMs) and orthologous DEGs are species specific

Next, we performed direct, cross-species comparisons using our FCG-derived rat dataset along with data from corresponding genotypes from our previous work in the FCG mouse model (Shin et al. 2025). We first used cross-species integration method SATURN to derive macrogenes from sets of similar genes across species (based on protein sequence similarity; **Sup. Table 9**). We then used the macrogenes to integrate data between species (**Fig. 4A-B**), create metacells (**Fig. 4C-D**), and perform differentially expressed macrogene (DEM) analysis (**Fig. 4E**).

**Figure 4.**
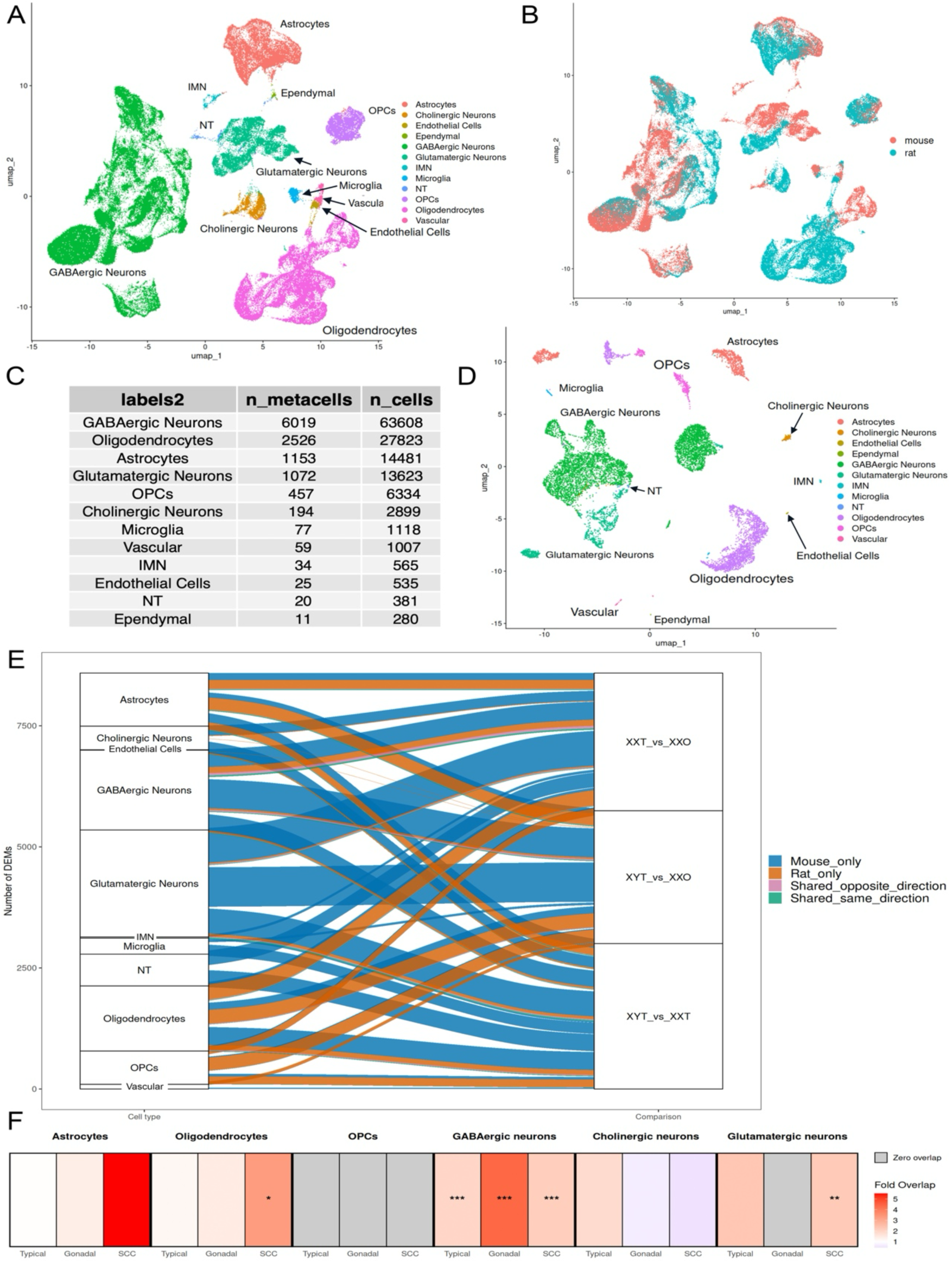
Differentially expressed macrogene (DEM) analysis after cross-species integration and metacell construction shows species-specific DEMs at individual macrogene level and limited overlaps in DEGs between species. (A) UMAP visualization of MS cell clusters colored by cell type after SATURN integration shows conservation of cell type identities between species. (B) UMAP visualization of MS cell clusters colored by species identity after SATURN integration shows successful cross-species integration and cell type homology where cells from FCGderived rats and FCG mice are within shared cell-type clusters. (C) Numbers of single nuclei and constructed metacells per cell type are proportionate. (D) UMAP visualization of MS cell clusters colored by cell type identity after metacell construction shows conservation of cell type identities. (E) River plot showing DEMs according to species origin reveals that the majority of DEMs are species-specific. Cross-species integration reveals conserved cell-type identity despite divergence in sex-biased expression of individual macrogenes between mouse and rat. (F) Direct comparison of sex-biased DEGs identified in rat and mouse shows significant overlaps limited to select cell type (e.g., GABAergic neurons) and sex factor (SCC) across species. Asterisks indicate overlap significance by Fisher’s exact test (* p0.05, **p<0.01, *** p<(0.001) Overall, limited DEM and orthologous DEG overlaps suggest divergence at individual gene expression level between mice and rats.

Results (**Fig. 4E**) show that the majority of DEMs are species-specific – suggesting that mice and rats seem to have diverged significantly in sex-biased expression of specific genes. Orthologous DEG overlap analysis (**Fig. 4F**) also showed limited overlap between species. Overlap was most significant across SCC comparisons and in GABAergic neurons (full statistics shown in **Sup. Tab. 10**). These limited gene level overlaps appear to contradict with higher level conservation in SCC dominance and network similarity between species. For example, in glutamatergic neurons, GRNs and key drivers from both rats (**section 2.5, Sup. Tab. 6**) and mice(Shin et al. 2025) are associated with mitochondrial and synapse functioning; these similarities in network function exist despite minimal DEG/DEM overlap. It is possible that downstream similarities in pathway and network sex differences may represent a case of convergent evolution – in which both rats and mice share similar downstream outcomes despite arriving at those outcomes through sex-biased expression of differing gene sets.

### Sex-biased DEGs are associated with Human Conditions and Traits

Next, we used Mergeomics to perform MSEA and identify significant disease associations in our DEGs for each sex factor and cell type using full summary statistics of 197 human genome wide association studies (GWAS) datasets available at GWAS catalog (select significant diseases in **Fig. 5A**, full statistical details in **Sup. Tab. 2**). Of these, 128 GWAS studies showed significant enrichment of disease-associated single-nucleotide polymorphisms (SNPs) in at least one cell type/sex-factor DEG set in the MS – implying potential roles for the sex-biased genes in human neurological, neuropsychiatric, and neurodegenerative disorders and related traits including cerebrovascular disease; PD, Cerebral Palsy (CP); developmental, eating, sleep, pain, hoarding, phobic, and panic disorders; cognitive function, memory, and intelligence; MDD; PTSD; Scz; neuroticism; ASD; headache and migraine; basal ganglia disease; AD; stroke; meningitis; anxiety; epilepsy; Tourette’s Syndrome; etc. As the sex-biased DEGs showing disease enrichment are expressed in other brain regions as well, these results do not directly imply that the MS is actually involved in the pathology of these conditions. However, it is interesting that many genes which are sexually dimorphic in the rat MS show enrichment for many human diseases that are sex biased (McCarthy et al. 2017).

**Figure 5.**
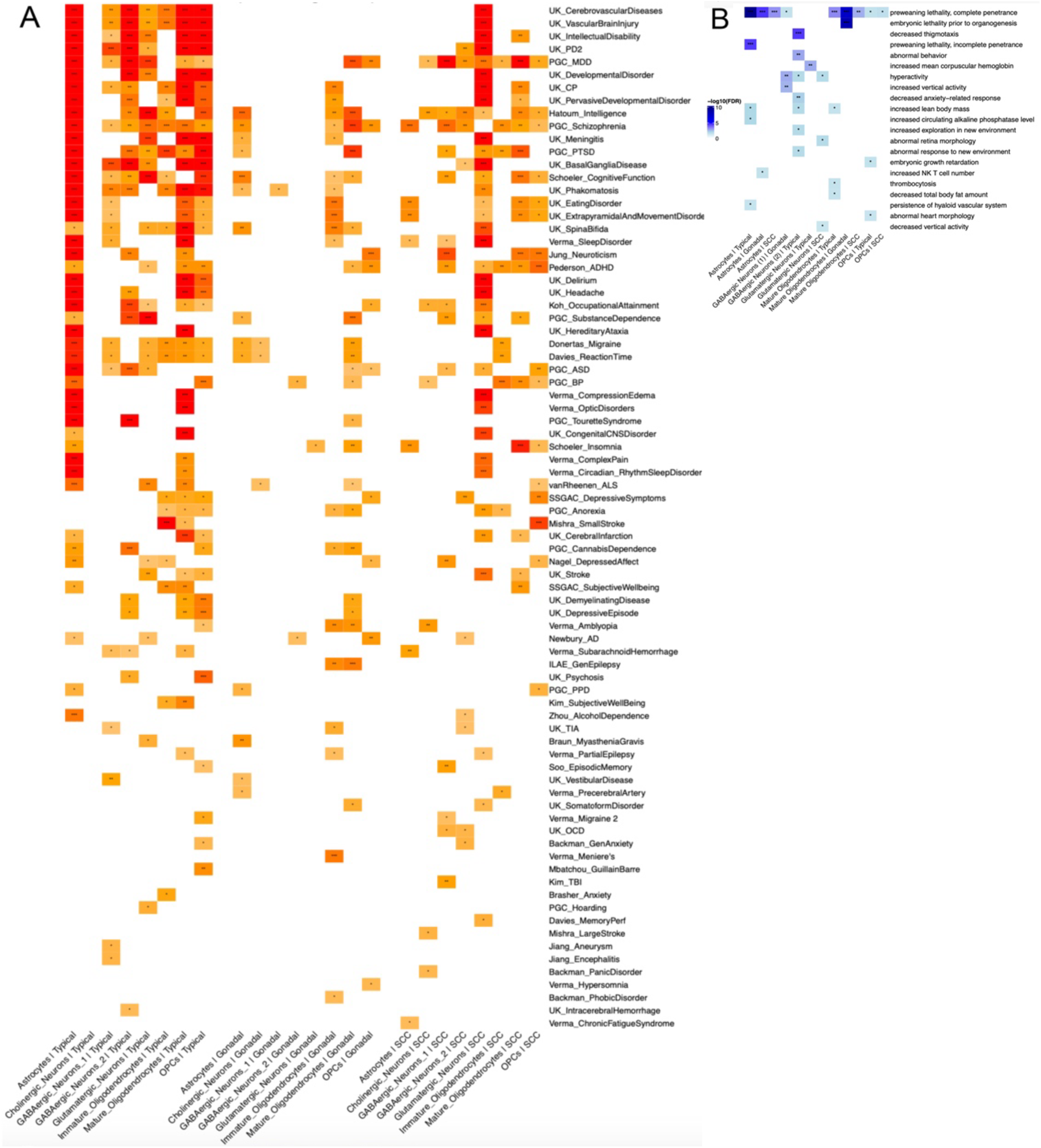
Mergeomics integration of 197 human GWAS summary statistics reveals enrichment of sex-biased MS DEGs from the FCG-derived rat with genes from numerous human neurological, neuropsychiatric, and neurodegenerative conditions and related traits, and integration with International Mouse Phenotyping Consortium (IMPC) data shows enrichment for various anxiety-related mouse phenotypes. Heatmap shows a sampling of significant human disease association results in the rat MS based on Mergeomics MSEA analysis where cell-typespecific DEGs across sex effects were used as input alongside mapped genes from human GWAS summary statistics targeting a range of neuropsychiatric, neurological, and neurodegenerative conditions and related traits (A). Cells are clustered by sex effect and cell type identities, color corresponds to degree of significance of enrichment based on -log10 transformed adjusted p-values, and asterisks denote specific significance thresholds (* p<0.05, ** p<0.01, *** p<0.001). Where multiple GWAS exist for one trait, the one with the most enrichment across cell types is shown. Results show the most associations between DEG sets belonging to cell types within the Typical comparison followed by those within the SCC comparison, and Typical Astrocytes show enrichment for 43 out of the 50 most significant (top) diseases and traits. These analyses identify widespread associations between sexbiased medial septum transcriptional programs and human neurological, psychiatric, and neurodegenerative traits. Heatmap shows the significant results of IMPC overexpression analysis wherein DEG clusters were assessed for enrichment with various mouse knockout phenotypes (B). Cells are clustered by sex effect and cell type identity, color corresponds to degree of significance of enrichment based on - log10 transformed adjusted p values, and asterisks denote specific significance thresholds (* p<0.05, ** p<0.01, *** p<0.001). Although some musculoskeletal, developmental phenotypes are likely not relevant to the MS, several phenotypes related to anxiety-like behavior support previous findings linking the MS to anxiety.

The present results contrast with our previous findings from the FCG mouse (select significant diseases in **Sup. Fig. 4, full statistical details in Sup. Tab. 2**), which we used here to run a Mergeomics analysis for cross-species comparison using matched XXO, XXT, and XYT groups(Shin et al. 2025). In the mouse, fewer (107 vs. 128) diseases and traits were significantly enriched compared to the rat. Also, while more enrichment was seen in the Typical and SCC DEGs in the rat, Gonadal DEGs in the mouse also showed strong disease enrichment. One similarity between species was the strong association of astrocyte DEGs, particularly Typical DEGs, with various diseases – suggesting a tight association between this cell type and various human pathologies.

### Enrichment of Sex-biased DEGs for Anxiety-like Behaviors in Gene Knockouts Mice

Finally, to infer potential causal effects of the DEG sets, we ran an enrichment analysis to compare our DEGs with gene knockout (KO) phenotypic associations curated in the International Mouse Phenotyping Consortium (IMPC) database (**Fig. 5B, Sup. Tab. 11**). Because the IMPC data is based on whole-body KOs, DEGs were enriched for various developmental, musculoskeletal phenotypes which likely are not relevant to the MS. However, results showed that Typical DEGs within GABAergic neurons (2) are enriched for KO phenotypes including decreased thigmotaxis, abnormal behavior, decreased anxiety-related response, abnormal response to new environment, hyperactivity, and increased exploration in new environment. It is interesting that many of these phenotypes are associated with anxiety-like behavior in mice since our MSEA analysis of human GWAS above also shows enrichment for anxiety disorders (**section 2.8**), and the MS has previously been linked to anxiety and stress (Wu et al. 2024).

## Discussion

Here, using a newly generated FCG-derived rat model, we document cell-type-specific sex differences in gene expression in the rat MS and show that SCC effects play a dominant role in the regulation of genes, pathways, and networks in the majority of cell types (**Fig. 7**). However, specific functional networks are influenced by either SCC, gonadal, or combined effects. Our results support the combined influence of these two sex-biasing factors in the Typical sex difference, and the affected genes are associated with various conditions in human GWAS and mouse KO phenotypes.

**Figure 6.**
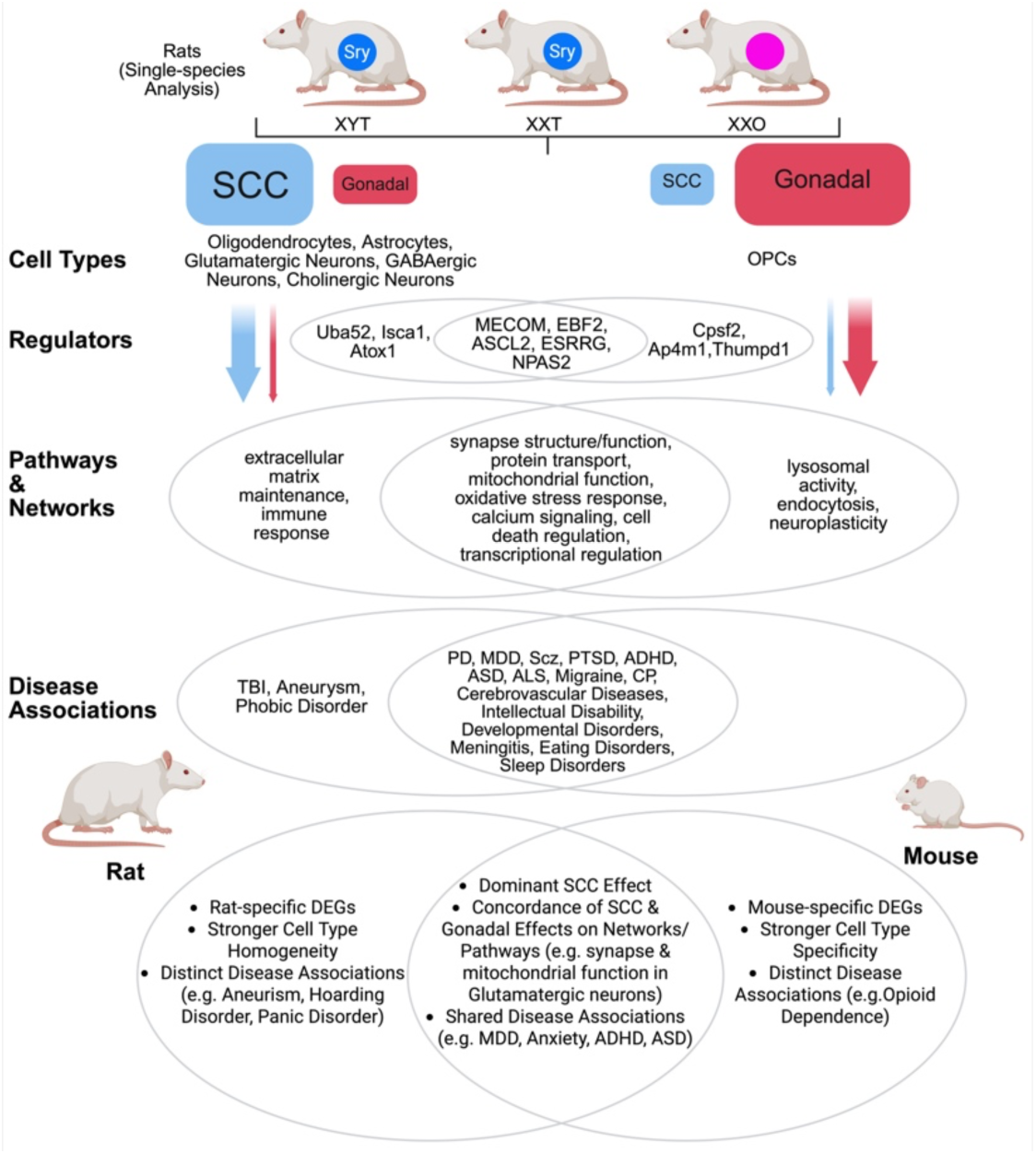
Summary of key findings from the FCG-derived rat model and rat vs mouse comparison. Our study of the FCG-derived rat model revealed dominant SCC regulation of gene expression within most cell types in the MS, likely through TFs and network key drivers which regulate downstream pathway and network functionality and disease associations – which all show dominant SCC effects in most cell types. Gonadal contributions are also important especially on the network level and in OPCs. Crossspecies analyses revealed similarities between FCG-derived rats and FCG mice including SCC effect dominance, concordance of SCC and Gonadal effects, and many shared disease associations. However, differences also exist; there were many species-specific DEGs, several species-specific disease associations, and differences related to celltype homogeneity or heterogeneity in the influences of SCC and Gonadal effects.

Our results agree with prior imaging-based work showing that sex-biased variations in MS volume were driven by SCC effects(Raznahan and Disteche 2021). Our results also align with previous cell morphology findings implicating SCC effects. First, Corre et al. demonstrated that the septal nuclei (including the MS) were larger in XXO and XXT mice compared to XYO and XYT (Corre et al. 2016). Then, Vousden et al. found no evidence that this pattern was changed in the adult MS when gonadal hormones were eliminated before puberty, despite a significant SCC × gonadal interaction effect in surrounding brain regions including the lateral septum(Vousden et al. 2018). Our findings extend prior work to further implicate SCC as the dominant sex-biasing factor associating with cell type-specific, sex-biased gene expression in the rat MS and provide molecular insights into potential mechanisms underlying the morphological and volumetric phenotypes observed earlier.

In our recent study of the mouse MS, Shin et al. explored the roles of SCC, gonadal, sex chromosome dosage, and interaction effects on gene expression across eight genotype groups and found evidence of strong SCC effects, which were largely concordant with gonadal effects(Shin et al. 2025). Importantly, all sex-biasing factors affected the expression of genes linked to human disease. Present results using matched genotype groups recapitulate these findings in both the rat and mouse and expand the list of human conditions linked to the MS in both species. Finally, they demonstrate the effectiveness of the novel FCG-derived rat model for studying sex-biasing effects with potential translational relevance to human pathology.

Comparing our present results with those from our recent work in the FCG mouse(Shin et al. 2025), both studies found similar cell types in the MS and showed that sex-biasing factors do not drive changes in cell type proportions across groups (**Fig. 6**). However, DEG clusters in the rat mainly showed concordance of Typical and SCC effects for individual cell types, but mouse DEG clusters seemed to discriminate based on cell type more than sex-biasing factor. Despite the differences, both the current rat and previous mouse studies showed concordance of SCC and gonadal effects on gene, pathway, and network regulation levels. GRNs were regulated by various sex-biasing factors in a cell-type-specific manner. Network functions showed some similarity across species but also key differences. For example, in glutamatergic neurons, mitochondrial and synapse activities are common to both species, but autophagy and viral/immune stimulus response are rat-specific.

Both rat and mouse DEGs, and many GRN key drivers, are associated with numerous human conditions and related traits, many of which were not previously linked to the MS (**Sup. Tab. 2**). These associations require further study. Additionally, involvement of key drivers in specific functional networks within the MS generates new mechanistic hypotheses for future investigation and highlights the MS potentially as a key pathological hub in a wide array of human conditions – many of which have not been previously considered in this region. More broadly, these findings highlight the importance of deliberate model selection, as mice and rats each offer distinct biological and experimental advantages. Mice are smaller, less expensive to maintain, more extensively inbred, and have historically benefited from a broader genetic toolkit. In contrast, rats possess larger brains that facilitate neuroanatomical, electrophysiological, and neurosurgical approaches; exhibit a richer repertoire of complex cognitive and social behaviors—including juvenile rough-and-tumble play; and have served as the principal model for decades of behavioral, pharmacological, and systems neuroscience research (Parker et al. 2014; Ellenbroek and Youn 2016; Carter et al. 2020; Pellis et al. 2023). These features make the rat particularly well suited for studies of SCC and gonadal sex, while our disease enrichment analyses further suggest that some human disorders may be better modeled in the FCG-derived rat (e.g. aneurysm, phobic disorder, and hoarding disorder) and others in the FCG mouse (e.g. opioid dependence).

In our matched-genotype, cross-species analysis, we found that most DEMs and orthologous DEGs are species-specific. We speculate that the many downstream similarities noted between rats and mice in terms of sex-biased regulation of pathway and network functions result from different gene expression patterns that independently evolve in each species but converge on similar overall functional effects. However, we note that his comparison was only made in the Sprague Dawley rat and MF1 mouse and needs to be validated in additional strains. Nevertheless, it highlights the importance of such similar downstream outcomes as both species likely derive benefit from them. The prospect that other mammals, such as humans, may have also evolved similar sexually dimorphic patterns in pathway and network regulation and function is worth exploring further. Finally, these findings underscore the importance of using multiple model organisms for translational work, as there are many species differences.

The current study is limited in the following ways. First, we do not attempt to distinguish between organizational vs. activational gonadal effects (Arnold 2009). Similarly, we do not attempt to parse the mechanisms underlying SCC effects such as the X vs. Y dosage effects as in our previous work in the mouse (Arnold 2009; Shin et al. 2025; Arnold et al. 2026). Secondly, the current study measures gonadal effects only in XX rats, and SCC effects only in rats with testes, in contrast to the mouse study, in which gonadal effects were measured in several sex chromosome genotypes and SCC effects were measured in mice with either type of gonad. To allow for meaningful cross-species comparisons, we used matched experimental groups in our analyses. We also implemented SATURN, a cross-species integration method, to directly compare DEMs across species with matched genotypes and followed up this analysis with overlap testing of orthologous DEGs from matched genotypes. Also, the Allen Institute’s MapMyCells tool used for cell type annotation does not include either the MS or MS-specific cell types. Thus, cell types found exclusively in the MS were likely mislabeled and named as cell types which are found elsewhere in the brain but have similar gene expression patterns. Thus, we re-grouped and re-named cell types according to their UMAP clustering patterns and then verified our cell type annotations with cell type-specific marker expression analysis based on markers found in the literature. Finally, Shin et al. used a different dissection method for mouse MS collection (in contrast to the more precise microdissection used in the current rat study) and thus may include tissues outside the MS (Shin et al. 2025).

In sum, the novel FCG-derived rat model recapitulated the dominance of SCC effects on gene expression regulation in the MS cell types previously observed in the FCG/SCT mouse model but also showed key differences. The findings support the FCG-derived rat as a useful addition to the repertoire of models of sex differences and suggest both divergent (on the gene expression level) and convergent (on the pathway and network level) evolution to be confirmed in additional animal models and humans. Our study highlights the importance of working with multiple model organisms since many of our findings were species-specific. Our findings connecting the gonadal and SCC-associated networks with human genetic studies offer potential molecular mechanisms of various sex-biased diseases linked to the MS which require functional validation. Our PharmOmics drug repositioning analysis represents one potential avenue of functional validation through network targeting using the predicted drug candidates which may prove useful for developing therapeutics for sex-biased human conditions linked to the MS. Thus, the work done here highlights the importance of the SCC in gene regulation in the MS and will likely drive future mechanistic and translational research of sex differences in MS-associated human disease.

## Methods

### FCG rats – breeding and genotyping

XY rats carrying an autosomal *Sry* transgene (XYTG) were generated at UCLA(Arnold et al. 2026). XYTG males from founder strain 208 were transferred to Georgia State University and mated with wild-type XX female Sprague Dawley rats (Crl: CD(SD) strain code 001; Charles River, Wilmington MA), producing four genotypes: XX rats with ovaries (XXO), XX rats with testes (XXT), XY rats with testes (XYT), and XYTG. XYO rats could not be generated as the Y chromosome in this model still carries a functional *Sry*. Therefore, we call this model “FCG-derived” instead of the FCG rat model. Offspring genotype was determined by PCR via tail snip biopsy. The Y chromosome was detected using primers targeting *Med14*, yielding a 560 bp PCR product for Med14X and a 172 bp product for Med14Y (forward: CCTCCAGACCCTATTACCAA; reverse: ATCACTGTCAAGGTTGCTTC). The *Sry* transgene was detected using two sets of primer pairs producing PCR products of 259 bp (180E22.1; forward: GCAAAGGTTATGACAGGGAAA; reverse: CCTCTTCGCTATTACGCCAG) or 294 bp (180E22.2; forward: TGTGGAATTGTGAGCGGATA; reverse: TCTGCTGTCCCCTGAGTTTT).

Rats were housed under a 12 h reverse light/dark cycle with unrestricted access to food (LabDiet 5001) and water. At postnatal day (p) 21, offspring were weaned into same-sex pairs in individually ventilated cages containing ALPHA-dri bedding and a standard enrichment tube. No more than 2 rats/genotype/litter were collected for this study. All experiments were approved by Georgia State University’s Institutional Animal Care and Use Committee and conducted in accordance with National Institutes of Health guidelines. Experimental design is in **Sup. Fig. 5**.

### MS dissection, nuclei isolation, and snRNA-seq library preparation

Adult rats (p78) were euthanized by rapid decapitation. Brains were removed, flash frozen in dry ice-cooled isopentane, and stored at −80°C until processing. Coronal sections were cut on a cryostat at −20°C, and tissue samples were collected from the brain surface using Rapid-Core sampling punches (Electron Microscopy Sciences). Micropunches targeting the MS were obtained using a 1.0mm diameter punch tool, with anatomical boundaries defined according to the Paxinos and Watson rat brain atlas (4th edition) beginning at 0.0mm medial-lateral, 1.0mm anterior-posterior, and -6.3mm dorsal-ventral to a depth of approximately 1mm.

Nuclei were isolated from MS samples from XXO, XXT, and XYT groups (n=6 rats/group totaling 18 rats across 3 genotypes, yielding 9 independent pools, with each pool containing MS from 2 rats) using a modified Dounce homogenization protocol. Tissues were placed on a chilled glass slide and manually minced with a sterile scalpel for approx. 1min until a homogeneous, paste-like consistency was achieved. Resulting tissue pulp was transferred to a pre-chilled Dounce homogenizer containing 300μL of chilled Lysis Buffer (487.5μL EZ Lysis Buffer [NUC101, Sigma-Aldrich] supplemented with 12.5μL RNase inhibitor). The slide was rinsed with an additional 200μL of Lysis Buffer into the homogenizer. Mechanical lysis was performed in two stages using a KIMBLE Dounce homogenizer. First, the suspension was processed with loose-fitting Pestle A for 1min, followed by 5min incubation on ice to facilitate chemical lysis. The homogenate was further processed with tight-fitting Pestle B for 30s to promote nuclear release. The final homogenate was filtered through a 40μm nylon cell strainer into a 50mL conical tube. To purify nuclei, the strainer was washed with 200μL of Debris Removal Buffer (200μL Debris Removal Reagent and 0.2μL Reducing Agent B from Chromium Nuclei Isolation kit). The filtered suspension was transferred to a 1.5mL low-binding microcentrifuge tube and centrifuged at 500g for 5min at 4°C with a reduced deceleration profile (brake set to 5). Supernatant was removed following centrifugation. The nuclei pellet was washed with Wash Buffer (1750μL 1X PBS, 200μL 10% BSA, and 50μL RNase inhibitor) and purified by centrifugation at 700g for 10min at 4°C followed by a wash in Wash Buffer.

Single nuclei suspensions were used to create Gel Bead-in-emulsions (GEMs) with the Chromium GEM-X Single Cell 3’ Reagent Kit v4 (10x Genomics, Pleasanton, California, USA). Library concentrations were determined using the Qubit Fluorometric Quantification method (Thermofisher Scientific, Waltham, MA, USA), and library quality was assessed using the Agilent TapeStation system (Agilent, Santa Clara, CA, USA). Pooled libraries were sequenced on the NovaSeq X Plus - 25B 2x150bp (Illumina, San Diego, CA, USA) in the UCLA Broad Stem Cell Research Center at ∼20k reads/cell.

### snRNA-seq pre-processing and quality control

10x Genomics’ Cell Ranger software (v9.0.0) was used to map FASTQ files to the refdata-gex-mRatBN7-2-2024-A reference genome. As snRNA-seq profiles nuclear transcripts including introns, intronic reads were not excluded. As Xist was not included in the reference genome, we added it following Twa et al with NCBI RefSeq annotation file (GCF_015227675.2) (Twa et al. 2025). Cell Ranger output files were processed via the CellBender package (v0.3.0) to remove signals derived from ambient RNA(Fleming et al. 2023). R package Seurat (v5.3.0) was used to load and combine ambient-RNA-corrected individual sample-wise gene expression matrices(Hao et al. 2024). Nuclei with <200 or >4,500 UMIs, <200 or >3,000 feature counts, or >1% mitochondrial percentage were filtered out. Remaining nuclei were annotated as predicted singlets or multiplets using DoubletFinder (v2.0.6) and scrubletR (v0.2.0) based on recommendations from benchmarking publications (Wolock et al. 2019; Xi and Li 2021; Kim et al. 2024), and multiplets called by either method were removed. This left an average of 7,836 nuclei per sample for downstream analysis. Quality control plots are shown in **Sup. Fig. 1.**

### Data normalization, batch correction, and cell clustering

We used Seurat functions “NormalizeData” to normalize gene expression matrices for individual samples, “RunPCA” for principal component analysis (PCA), and “RunHarmony” to correct for batch effects with the grouping variable set to “batch.” Batch refers to sample preparation date prior to snRNA-seq. All samples were sequenced simultaneously. The K-Nearest Neighbors and Louvain algorithms were implemented using Seurat’s “FindNeighbors” and “FindCluster” functions to cluster nuclei with similar expression patterns. We performed Uniform Manifold Approximation Projection (UMAP) to visualize clusters in 2D space using the top 40 PCA dimensions.

### Cell type annotations

Due to the lack of rat cell type references, we used mouse single cell references to infer cell types in the FCG-derived rat MS. Cell type annotation was performed using the Allen Institute’s MapMyCells tool (RRID:SCR_024672) (Daniel et al. 2026) using hierarchical mapping to the 10x Whole Mouse Brain (CCN20230722) reference taxonomy after conversion of rat genes into mouse orthologs using R package GeneOrthology (v0.1.0) (Science 2024). Cells with <95% mapping certainty were removed as well as cell types with <10 cells. Additionally, those cell types not found in the MS and with <100 cells were removed (see below). This left 70,512 single nuclei for downstream analysis. As the MS has not been explicitly profiled in the Allen Brain Atlas and thus not labeled in the MapMyCells anatomical annotations, cell type names can be from non-MS brain regions whose cell types share gene expression similarity with MS cells (e.g., MS cells mapped to MapMyCells subclass “064 STR-PAL Chst9 Gaba” are not from the striatum or pallidum but share gene expression similarity to cells found there). To avoid confusion, we modified cell type names according to UMAP clustering patterns (**Sup. Tab. 1**; e.g. we renamed “064 STR-PAL Chst9 Gaba” to “GABAergic neurons (2)”). Finally, the “Oligodendrocyte” cell type was split into “Mature Oligodendrocytes” and “Immature Oligodendrocytes” based on their expression of specific markers, including *Bcas1* and *Gpr17* (immature oligodendrocytes), *Cspg4* (OPCs), and *Cldn11* (oligodendrocytes) (Chen et al. 2009; Fumagalli et al. 2011; Kaji et al. 2020; Beiter et al. 2022; Gjervan et al. 2023). All cell types were verified based on expression of canonical cell type markers: *GFAP* (astrocytes), *Ngfr* (cholinergic neurons), *Gad1* (GABAergic neurons), *Slc17a7* and *Slc17a6* (glutamatergic neurons), *P2ry12* (microglia), *Cldn11* (oligodendrocytes), *Gpr17* and *Bcas1* (immature oligodendrocytes), *Cspg4* (OPCs), *Cd34* (endothelial cells), and *Kcnj8* and *Ogn* (vascular cells) (Shanahan et al. 1997; Chen et al. 2009; Fumagalli et al. 2011; Kaji et al. 2020; Ando et al. 2022; Beiter et al. 2022; Langlieb et al. 2023; Radu et al. 2023; Yao et al. 2023; Kuhn et al. 2024; Mei et al. 2024).

### Genotype Validation Using snRNAseq Data

To confirm sample genotypes, we assessed expression of sex-linked genes *Uty* and *Xist* post-sequencing and did not identify any mis-genotyped samples.

### Cell Type Proportion Analysis

Cell type proportions were calculated for each sample individually as the number of nuclei per cell type divided by the number of nuclei per sample. We used the propellor method from R package speckle (v1.10.0) (Phipson et al. 2022) to detect significant differences in cell type compositions between groups via an empirical Bayes moderated one-way ANOVA on logit-transformed cell-type proportions.

### Generation of Metacells & Identification of Differentially Expressed Genes (DEGs)

Benchmarking studies recommend using a metacell-based approach for DEG identification (Bilous et al. 2022; Bilous et al. 2024). This method groups transcriptionally similar cells into a single representative cell state – a “metacell”. Metacells are constructed within a given cell type and sample to better preserve biological variation and reduce data sparsity and biases stemming from pseudoreplication. This method has already been applied successfully in various settings for DEG identification (Giladi et al. 2018; Sebe-Pedros et al. 2018; Liu et al. 2022; Kirschenbaum et al. 2024; O’Leary and Zheng 2024). Here, metacells were created using R package MetaCell (v0.3.7) and the following parameters: k = 10, reduction = “pca,” dims = 1:40, layer = “counts”, mode = “sum”, min cells = 10, max shared = 1, and ∼10 cells/metacell (Baran et al. 2019). Metacell proportions per sample and cell type reflect original cell type abundances.

To determine the contributions of typical sex difference and individual sex factors to gene expression in each cell type, pairwise linear models were fitted across all relevant samples using R package Limma (v3.66.0) (Ritchie et al. 2015) to identify significant DEGs in each between-groups comparison (Typical or Wild Type: XYT vs. XXO, Gonadal: XXT vs. XXO, and SCC: XYT vs. XXT). In this design, the gonadal effects are tested only in XX rats, and SCC effects only in rats with testes. All linear models included batch as a covariate to account for technical batch effects. “Batch” refers to sample preparation date prior to snRNAseq. For each comparison, the linear model used was: *Υ (gene expression) = β_0_ + β_1_Group + β_2_Batch + ε* Differential gene expression was calculated from these models. The false discovery rate (FDR) was calculated using the Benjamini-Hochberg (BH) approach. DEGs with FDR <0.05 and log2 fold change (log2FC) >0.1 were considered significantly differentially expressed. The Wilcoxon test with BH correction was used to compare *Crhbp* levels across groups.

### DEG Clustering and Pathway, Transcription Factor Enrichment Analysis

Because of the large numbers of DEG sets across cell types and sex factors, we identified DEG sets (clusters) sharing similar expression change patterns between sex effects and cell types to simplify interpretation. The log2FC values of DEGs were isolated for each cell type and sex effect pair and then correlated with all others via the Pearson method. All DEGs, regardless of significance, were used with a median of 16,496 genes/cell type/sex effect. We observed clear separation of clusters based on dendrogram patterning and biological interpretability. Significant DEGs within each cluster were converted to mouse orthologs using R package biomaRt(Durinck et al. 2005; Durinck et al. 2009) (v2.66.1) and used as input for (1) pathway enrichment analysis using R package clusterProfiler (v4.18.4) (Yu et al. 2012) and gene sets from the GO Biological Process (GO:BP) database(Ashburner et al. 2000; Gene Ontology 2026) and (2) TF enrichment analysis using enrichr (Chen et al. 2013; Kuleshov et al. 2016; Xie et al. 2021) and the PWM Mouse 2025 database (Rauluseviciute et al. 2024). Pathways or TFs at FDR <5% were considered significantly overrepresented. Redundant pathways with highly overlapping genes were identified using R package rrvgo (v 1.22.0) via the Jaccard index coefficient.

### Gene Regulatory Network (GRN) Construction

For each cell type, global gene regulatory networks (GRNs) were created using Single Cell Integrative Gene Regulatory Network Inference (SCING) and cell type specific gene count matrices (Littman et al. 2023). Gradient boosting regressors were trained on each gene to form directed edges from potential upstream regulators. Then the weighted Key Driver Analysis (wKDA) module from R package Mergeomics (c1.38.0) was used to overlay cell type specific sex factor DEG sets onto the constructed networks (Shu et al. 2016; Ding et al. 2021). This enabled identification of key drivers whose neighborhood subnetworks were significantly enriched for sex-biased DEGs at FDR <5%.

### PharmOmics Drug Repositioning

Drug repositioning analysis was performed for each GRN using PharmOmics (Chen et al. 2022) to identify drug molecules that could modulate the sex-biased subnetworks. The app2 module was used for drug repositioning based on the connectivity between PharmOmics drug signatures and key drivers from the wKDA analysis within SCING-derived GRNs. Drugs with z-score p-values with BH correction <0.05 were considered significant. Jaccard scores indicate the amount of overlap between input genes and genes on which PharmOmics drugs are expected to act. Jaccard scores were summed across cell types for each significant drug to prioritize drugs acting on sex-biased genes across cell types.

### Disease Enrichment Analysis

We explored the relevance of cell-type-specific sex factor DEGs to human diseases. We first collected 197 summary statistics from human GWAS for a range of neuropsychiatric, neurodegenerative, and neurological traits and diseases available in GWAS catalog (**Sup. Tab. 2**)(Cerezo et al. 2025). GWAS summary statistics were included for studies published in 2020 or later, with sample size of at least 100,000, and/or if they were included in our previous analysis for FCG mouse MS by Shin et al. (Shin et al. 2025) Then, using the Marker Dependency Filtering (MDF) module from Mergeomics, GWAS single nucleotide polymorphisms (SNPs) were linked to genes via 50kb distance mapping and corrected for linkage disequilibrium with cutoff at r^2^>0.5. Significant sex-biased rat DEGs were converted into human orthologs using R package biomaRt. We then used the Marker Set Enrichment Analysis (MSEA) function in Mergeomics to determine if sex factor DEGs show stronger GWAS association than would be expected by random chance. This was done by comparing disease association p-values of SNPs mapped to our DEG sets to those of the SNPs mapped to 10,000 sets of random genes using a χ2-like statistic. Permutation-based p-values were calculated, and the BH method was used to derive FDR. Significance was set to FDR <5%. We repeated this analysis on DEGs from our previous work in the FCG mouse using matched genotypes (XXO, XXT, and XYT) to make direct comparisons across species (Shin et al. 2025).

### Gene Knockout Phenotype Enrichment Analysis

To infer potential causal effects of the sex-biased DEG sets, enrichment of DEGs for mouse phenotypes in genetic knockout models was evaluated via R package clusterProfiler (v4.18.4) and genes from the International Mouse Phenotyping Consortium (IMCP) (www.mousephenotype.org) after converting rat DEGs to mouse orthologs (Dickinson et al. 2016; Wilson et al. 2026). Phenotypes at FDR <5% were deemed significantly overrepresented.

### Cross-species Integration and Comparison

We next performed direct cross-species comparisons using SATURN. This method uses protein sequence similarity and machine learning to construct “macrogenes” which include sets of similar genes from one or more species. Macrogenes are then embedded into individual species and used for differential expression(Rosen et al. 2024). Here, we integrated data from all rat groups (XXO, XXT, and XYT) with corresponding groups from the FCG mouse. We then constructed metacells and performed differentially expressed macrogene (DEM) analysis within species in the same manner as DEG analysis before (**section 1.8**) to determine whether results were similar across species.

Additionally, we performed an overlap analysis between rat DEGs and orthologous mouse DEGs matched by cell type and sex factor from genotype-matched mouse data published by Shin et al (Shin et al. 2025). BiomaRt was used to determine DEG orthologs, and a Fisher’s exact test with BH correction was used to determine whether DEG overlap between species was significant within each cell type

## Software Availability

The code used to analyze the data are deposited to GitHub at https://github.com/XiaYangLabOrg/FCG-Rat-MS

## Data Availability

The snRNAseq data generated in the current study have been deposited to Gene Expression Omnibus with accession number GSE349124.

## Supplemental Information

Document S1. Supplementary Figures 1-5 and Supplementary Table 1 with titles and legends.

Supplementary_Table_2. Full list of GWAS traits/diseases with citations and full results.

Supplementary_Table_3. Full list of DEGs.

Supplementary_Table_4. Full results of cluster pathway analysis.

Supplementary_Table_5. Full results of cluster TF analysis.

Supplementary_Table_6. Full list of key drivers and their known functions with citations.

Supplementary_Table_7. Full results of PharmOmics analysis.

Supplementary_Table_8. Top 25 drugs resulting from PharmOmics analysis.

Supplementary_Table_9. Full results of SATURN DEM analysis.

Supplementary_Table_10. Full results of rat/mouse orthology overlap analysis.

Supplementary_Table_11. Full results of IMPC overlap analysis.

## Acknowledgments

This work was supported by the National Institutes of Health (R21 MH129020, R01 DA056534) and the National Science Foundation (IOS-2313253).

## Author Contributions

VT performed computational analyses and wrote the manuscript. GZ performed sample preparation and computational analysis. EH performed animal husbandry and tissue collection and validation. IA performed sample preparation. JS created the original data analysis pipeline. MR, ET, DR, and RW participated in result interpretation and manuscript writing. MD, VH, AG, and AA contributed to the development of the FCG-derived rat model used in the current study. AA, DB, and XY performed supervisory and experimental design roles. All authors contributed to the manuscript.

